# Lightsheet microscopy integrates single-cell optical visco-elastography and fluorescence cytometry of 3D live tissues

**DOI:** 10.1101/2024.04.20.590392

**Authors:** Yuji Tomizawa, Khadija H. Wali, Manav Surti, Yasir Suhail, Kshitiz, Kazunori Hoshino

**Affiliations:** Department of Biomedical Engineering, University of Connecticut, CT; Department of Biomedical Engineering, University of Connecticut Health, Farmington, CT; Systems Biology Institute, Yale University, West Haven, CT

## Abstract

Most common cytometry methods, including flow cytometry, observe suspended or fixed cells and cannot evaluate their structural roles in 3D tissues. However, cellular physical interactions are critical in physiological, developmental, and pathological processes. Here, we present a novel optical visco-elastography that characterizes single-cellular physical interactions by applying in-situ micro-mechanical perturbation to live microtissues under 3D lightsheet microscopy. The 4D digital image correlation (DIC) analysis of ∼20,000 nodes tracked the compressive deformation of 3D tissues containing ∼500 cells. The computational 3D image segmentation allowed cell-by-cell qualitative observation and statistical analysis, directly correlating multi-channel fluorescence and viscoelasticity. To represent epithelia-stroma interactions, we used a 3D organoid model of maternal-fetal interface and visualized solid-like, well-aligned displacement and liquid-like random motion between individual cells. The statistical analysis through our unique cytometry confirmed that endometrial stromal fibroblasts stiffen in response to decidualization. Moreover, we demonstrated in the 3D model that interaction with placental extravillous trophoblasts partially reverses the attained stiffness, which was supported by the gene expression analysis. Placentation shares critical cellular and molecular significance with various fundamental biological events such as cancer metastasis, wound healing, and gastrulation. Our analysis confirmed existing beliefs and discovered new insights, proving the broad applicability of our method.

## Introduction

Digital image correlation (DIC) is emerging for materials characterization, inspection, and testing [1,2]. Light microscopy-based displacement and deformation analysis, or optical elastography, have been successful in biosample characterization. Optical Coherence Microscopy (OCT) based elastography is closely related to our study and has succeeded in microscopic tissue structural characterization [3,4]. Traction force microscopy (TFM) [5-7] allows for the measurement of cellular contractility and is used to study cells in a 2D monolayer culture. However, there is still an unmet need for rapid assaying to investigate the correlation between structural and molecular characteristics in 3D tissues at the single cellular level. Current methods can not provide high-resolution 3D structural analysis directly linked to fluorescence assays. Flow cytometry tools do not examine heterotypic cellular interactions that occur inherently in 3D tissues. Brillouin imaging (BI), used for biological tissue characterization [8-10], is a point scan method and requires a long acquisition time to image 3D tissues.

Analysis of structural cellular properties in 3D tissues has been of great interest in biology. Cellular and tissue viscoelastic characteristics are crucial for development, regeneration, and tumor invasion. Notable processes include progression of development through the interaction of dermal layers in gastrulation [11,12], sprouting of blood vessels [13], invasion at the maternal-fetal interface [14], and desmoplastic reaction to cancer [15-17]. In particular, the transitions between fluid-like (viscous) and solid-like (elastic) behaviors permit cells to undergo different modes of single-cell and collective-cell migration [18]. Epithelial-mesenchymal transition (EMT) and mesenchymal-epithelial transition (MET) play critical roles in embryonic development, tumorigenic process [19], and wound healing [20]. Those processes involve transitions in the viscoelastic properties of the cells, which are critical to the functional outcome of these processes. A recent pioneering study showed the transition from a fluid-like state to a solid-like state in embryonic tissues [21], where the analysis was made by injecting magnetically responsive ferrofluid, which gives information about a relatively small neighborhood around the droplet [22] rather than a whole tissue.

Here we take an optical imaging approach, using lightsheet microscopy [23,24] and 4D DIC analysis to combine the benefits of fluorescence-based cytometry and single cellular visco-elastography, linking molecular assays and image analysis-based structural characterization. We have integrated a miniature mechanical perturbation device that works under a high-end light-sheet microscope (LSM). LSM is advantageous for the volumetric imaging of 3D tissues, allows for high resolution, high throughput, and multi-channel fluorescence imaging, and is suitable for investigating complex cellular and molecular interactions in the micro 3D environment [25-27]. Our unique method of volumetric DIC analysis examines cellular displacements and strain distribution induced by mechanical perturbation, revealing the viscoelastic structural interaction between single cells. We evaluated our method’s efficacy by studying a 3D organoid model of maternal-fetal interface in human placentation. In humans, decidualization involves a notable transformation of maternal endometrial stromal fibroblasts (ESFs), marked by increased contractile force generation, elevated matrix deposition, and enhanced cellular secretion, in preparation for placental trophoblast implantation and invasion [28,29]. We examined how cellular transitions affect the 3D tissue properties and verified that ESFs became stiffer when differentiated (dESFs). Our study also quantified that extravillous trophoblasts can partially alleviate dESF stiffness. Our innovative 4D DIC-based structural analysis enabled the observation of tissue viscoelasticity at the cellular level. Placentation shares critical cellular and molecular significance with fundamental biological events such as cancer metastasis, wound healing, and gastrulation. Our approach, thus, offers a powerful means to investigate viscoelastic dynamics at tissue interfaces, with broad relevance in both physiological and pathological spatial transitions.

## Results

### 4D Digital Image Correlation (DIC)

We built the image acquisition system on the high-end Zeiss Lightsheet7 microscope (**Figure 1 (a) top**). Our innovation integrates micro-mechanical perturbation and synchronized 4D imaging to record the compressive strain applied to the specimen. **Figure 1(a) bottom** shows a photograph of a microtissue sample in agarose on the compression table. A miniature stepper motor displaces the table by ∼5 μm for each compression step. As the table moves and applies stepwise compression to the agarose, the microscope acquires volumetric images of sample deformation. Two laser light sources (488 nm and 561 nm) and dual illumination objectives (10x NA 0.2) provide light sheet excitation from both sides. An imaging objective (20x NA 1.0), dichroic filters, and two sCMOS cameras (PCO.edge 4.2) acquired simultaneous two-channel fluorescence images. The sample deformation in response to compression is a time-dependent viscoelastic process. After each stage displacement, we applied ∼10 sec (typ.) of wait time before image acquisition. With the time constant of a spheroid compression reported to be less than 5 sec [30], we assume the sample is in mechanical equilibrium when imaged. For each compression step, we acquired the 3D volume (1,600 (x) × 1,600 (y) × ∼360 (typ.)) at a rate of 100 ms/ z layer (50 ms for each of left and right excitation), resulting in ∼60 sec/step including the wait time. The total image acquisition time for the entire micro-compression process (7 indentation steps) is thus typically ∼7 mins. **Figure 1(b) top** shows an acquired indentation sequence of a 3D organoid model of maternal-fetal interface with placental extravillous trophoblasts (EVTs) and endometrial stromal fibroblasts (ESFs). We built a custom 4D digital image correlation (DIC) program on MATLAB to conduct the spatially-resolved deformation analysis (see **Figure 1(b) bottom**). We acquired images for the hemispherical half of an organoid closer to the microscope objective as the region of interest. We then evenly distributed nodal points on the segment boundaries and inside the segments with ∼10 pixel (∼4 μm) distances, which resulted in a total of ∼20,000 nodal points. For the 4D digital image correlation, we adapted the algorithm reported in [31] and tracked the 3D coordinates of all nodes along the indentation steps. The DIC program used single-channel, grayscale images combining red and green fluorescence signals for node tracking.

**Figure 1.**
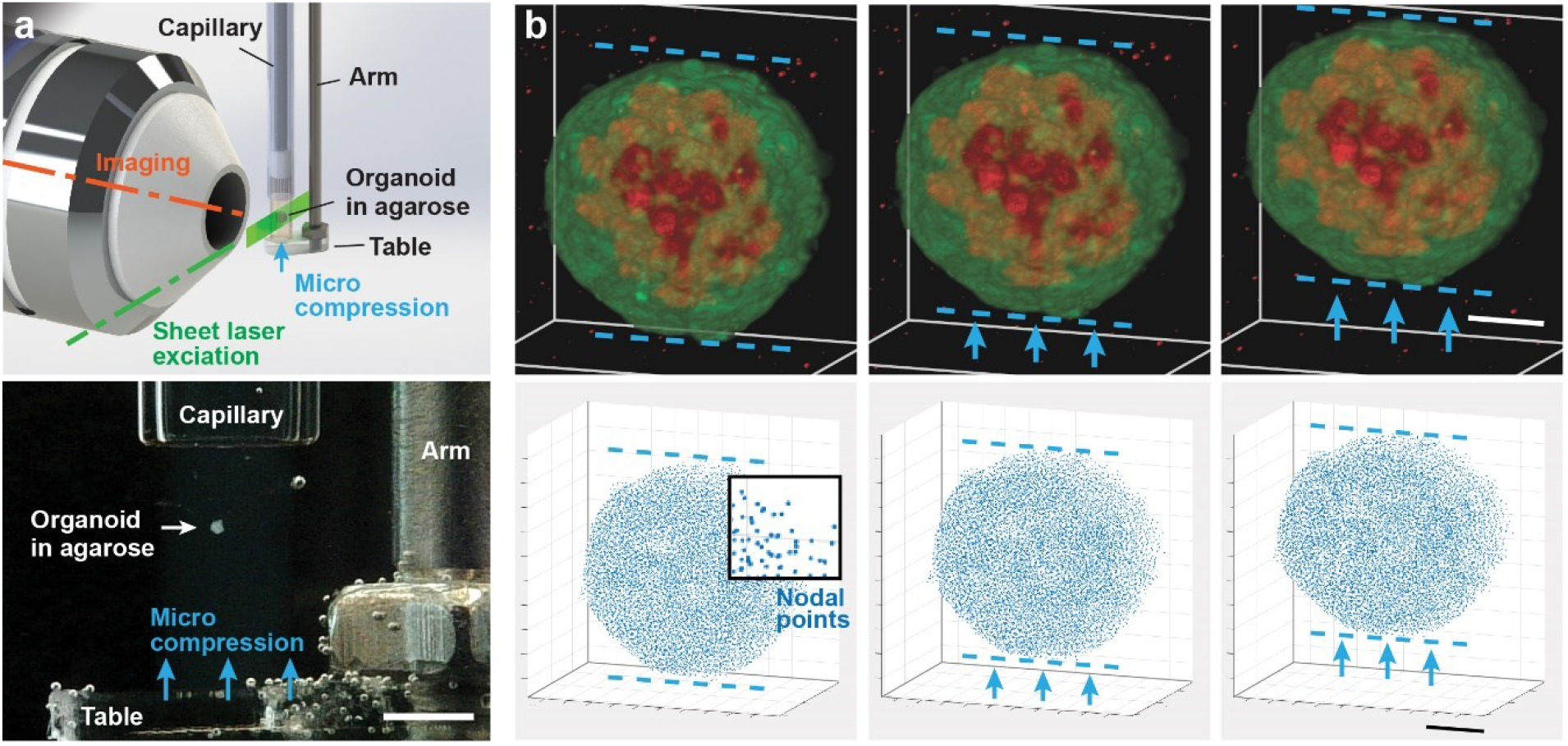
**(a)** The light-sheet microscopy integrated with the micromechanical perturbation device. An overview illustration (top) and an experimental photograph (bottom, scale bar = 1 mm) show the device applying compressive strain to the sample embedded in an agarose pillar. **(b)** A compression sequence acquired by our microscope. We imaged a 3D organoid composed of extravillous trophoblasts (green), and endometrial stromal fibroblasts (red) (top, scale bar = 50 μm). The 4D digital image correlation (DIC) algorithm tracked the features of ∼20,000 nodal points to obtain spatially-resolved displacements in 3D (buttom, scale bar = 50 μm).

### Single-cellular structural analysis revealed solid-like and liquid-like cell-cell interactions

The combination of digital image segmentation and 4D nodal tracking enabled the analysis of single-cell characteristics and interactions. We built a custom algorithm, modified from [32], to separate the imaged region into 500-600 segments representing single cells, which permits indexing and analysis of individual cells (see **Supplementary Information** and **Supplementary Figure 1** for details). **Figure 2(a)** shows an example of the cellular displacement distribution under compressive stress. We found the cellular displacement vectors by averaging those of nodal points included in each cell. Many cells were correlatively displaced along with neighboring cells as expected in an elastic body (see **Figure 2(b)** top marked ‘aligned’). On the other hand, we also observed regions with liquid-like deformation, where displacement vectors show an eddy-like pattern with backward or rotational motion (see **Figure 2(b)** middle and bottom). The shear stress intensity in fluid mechanics indicates the randomness of moving finite-sized fluid particles in a turbulent flow. Using a similar approach, we evaluated cellular shearing motion in the 3D organoid by the von Mises strain, which takes shear strain into account. We computed the strain by creating a mesh with 100,000-200,000 tetrahedral elements from the ∼20,000 nodal points tracked by the 4D DIC. Note that the number of tetrahedral elements is usually larger than the number of nodal points. Each element belongs to one of the 500-600 segments, representing single cells, forming the 3D organoid model. Each segment typically contained a few hundred elements. We calculated the segmental strain tensor from the volumetric average of elemental strains. **Figure 2(c)** shows the von Mises strain map calculated for the same sample. The local areas with larger von Mises strain (i.e., orange-red regions) indicate more intense distortion energy than surrounding regions, suggesting more liquid-like behaviors.

**Figure 2.**
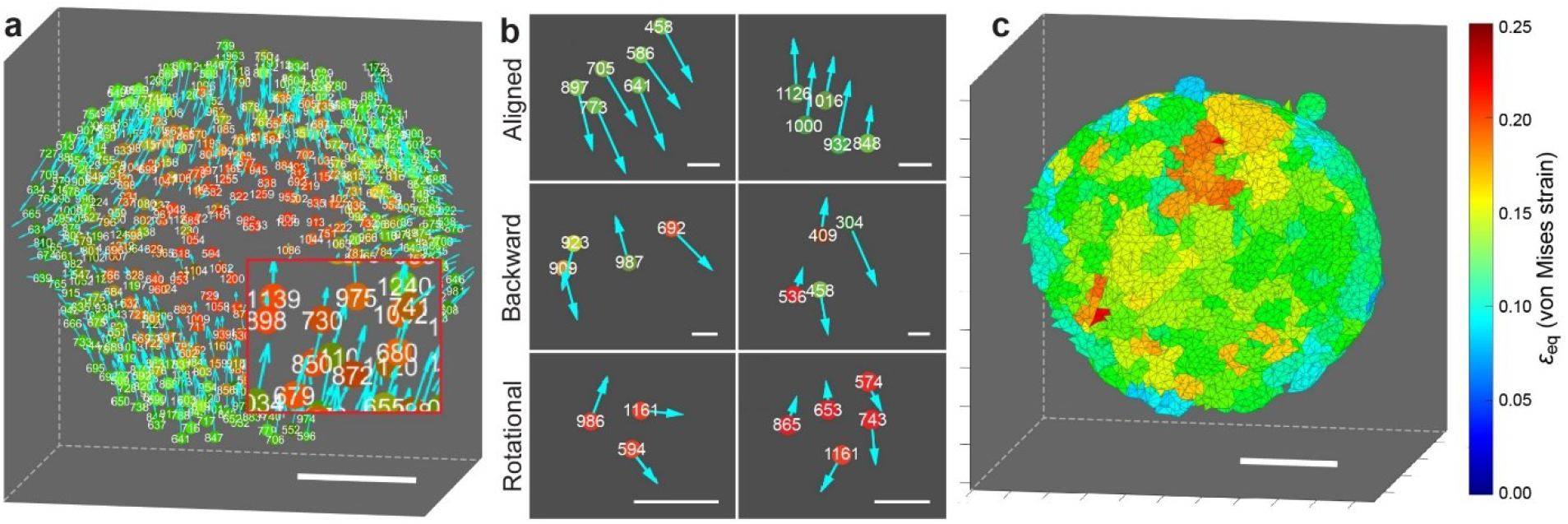
**(a)** Our analysis segments, indexes, and tracks individual cells to visualize the displacement field under compression. Scale bar = 50 μm. **(b)** We revealed different local patterns in cell-cell interaction, including solid-like well-aligned deformation (top), fluid-like local backward shearing (middle), and rotational (bottom) motion. Scale bars = 20 μm. **(c)** The displacement tracking of ∼20,000 nodal points allows for von Mises strain mapping, indicating the randomness in local deformation. Scale bar = 50 μm.

### 3D composite organoid model of maternal-fetal (endometrial-placental) interface

The maternal-fetal interface in mammals demonstrates a unique juxtaposition of tissues with different genetic backgrounds, which has given rise to an evolutionary phenomenon called “genetic conflict [33].” Among hemochorial placental mammals, such as great apes, which include humans, and rodents, placental trophoblasts of fetal origin invade deeply into the maternal endometrium [34,35]. Among humans, in anticipation of implantation, the maternal endometrial stromal fibroblasts (ESFs) differentiate into decidualized ESFs (dESFs), a process evolutionarily derived from myofibroblast-like transformation [36]. Decidualization, among many other changes, greatly increases the contractile force generation and collagen deposition by ESFs [37,38]. It was shown that recently evolved extravillous trophoblasts (EVT) in the placenta of great apes can partially reverse the decidual contractile force generation, promoting their own invasion into the maternal endometrium [39]. EVTs are characterized by aggressive invasion into the maternal tissues. We created a composite organoid model of the placental-decidua interface and used it to evaluate the efficacy of measuring cellular viscoelastic properties at the 3D tissue interface. Decidualization is both an adaptive response to provide sustenance to the placenta, as well as an evolved response to limit excessive invasion by trophoblasts. Decidualization has evolved from the fibroblast activation response, which is likely activated owing to the injury caused by implantation in the endometrial luminal epithelium [36,40]. A 2D model analysis has demonstrated that decidualization of ESFs significantly increases their contractile force generation [39], strengthening their resistance to trophoblast invasion. It was also shown that co-culture with HTR8, a cell line derived from EVTs, reversed the force generation capability of decidualized ESFs (dESFs). We used RNAseq data for human ESFs, dESFs, and dESFs treated with conditioned medium from HTR8 (dESFs^cond^). To ensure that the withdrawal of decidual stimulus did not result in the dedifferentiation of dESFs, we added progesterone (MPA) in the HTR8 conditioned medium.

Figure 3. shows the characteristics of the three samples we used in this study. We prepared organoids containing human endometrial stromal fibroblasts (ESFs) and placental extravillous trophoblast HTR8 cells. We tested three conditions of stromal fibroblasts, namely, undifferentiated (ESF), decidualized (dESF), EVT conditioned decidualized (dESF^cond^) fibroblasts (see **Supplementary Information** for the details). The organoid diameters ranged between 160-180 μm. We found that for all three cases (ESF, dESF, and dESF^cond^), self-organization of a mixed ESF-HTR8 ensemble resulted in a layer of trophoblast cells (green) enveloping the fibroblast cells (red), suggesting the surface or interfacial tension [41] of the trophoblast cells was lower than the fibroblast cells. The structural and detailed gene expression analyses summarized in the figure supported the myofibroblast activation in decidualization, and its reversal by paracrine signals from EVTs. **Figure 3(a)** shows the averaged structural characteristics of ESFs. We calculated the von Mises strain of the core regions (50% of the total volume) to compare the fibroblast stiffness. Indeed, dESFs showed significantly lower levels of strain, which was partially and significantly restored in dESF^cond^ after treatment with the EVT conditioned medium. Gene expression analysis conincidentaly supported the structural transition between the three types. Principal component analysis (PCA) for ESFs (**Figure 3(b)**) confirmed well-separated clusters for samples from all three conditions. We noticed that the changes induced by decidualization were nearly reversed in the PC2 axis, suggesting that HTR8 conditioning itself could partially reverse gene expression changes caused by decidualization. Of the total 1794 genes that had increased significantly from ESF to dESF, 1116 were reversed by HTR8 conditioning. Similarly, of the 1946 genes that were significantly decreased by decidualization, 1389 genes were significantly increased by HTR8 conditioning, confirming that EVTs can reverse the decidualization of ESFs. We then asked if the conditioned medium from HTR8 had any effect on gene-sets associated with cellular contractility, actomyosin assembly, and other ontologies associated with cellular force generation (**Figure 3(c)**). Overall, for most of the key contractility-related gene ontologies (GOs), decidualization resulted in significant activation, while conditioned medium reversed the trend. Notable among these GOs included muscle-contraction, myotube-differentiation, various ontologies associated with contractile actin filament assembly, as well as cell adherens junctions. Interestingly, we found that HTR8 conditioning resulted in increased activation of myosin-V-binding, as well as GOs associated with cell-cell adherens junction. The unconventional myosin V, a tubulin associated motor protein, is not involved in cellular contractility, but has the role of cargo transport in yeast and mammalian cells [42,43]. Overall, our data confirmed that the HTR8 could reverse the decidualization-induced activation of pathways involved in increased cellular force generation and cell-matrix interactions.

**Figure 3.**
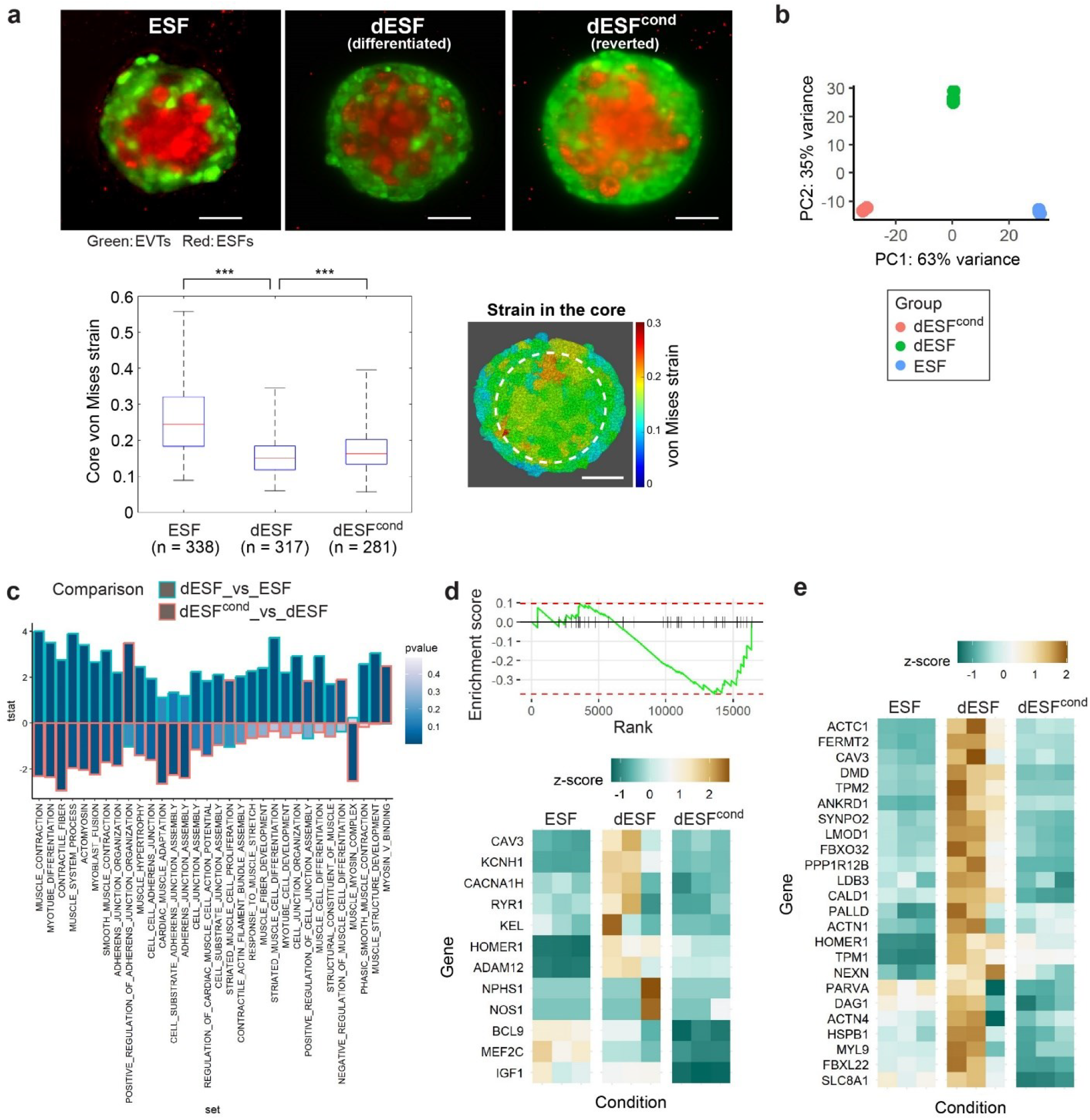
Characterization of undifferentiated (ESF), decidualized (dESF), and conditioned decidualized (dESF^cond^) fibroblasts we used in this study. The structural and gene expression analyses coincidently demonstrate the viscoelastic transitions between the three conditions. **(a)** For all three conditions, fibroblast cells occupied the core regions. Decidualized dESFs show smaller von Mises strains than ESFs, while dESFs^cond^ partially reversed the transition back and showed large strains. Scale bars = 50 μm. **(b)** Principal component analysis (PCA) plot showing global gene expression (n = 3 biological replicates) demonstrated a similar trend in the PC2 axis. **(c)** Relative activation of gene ontologies (GOs) associated with myofibroblast activation. Decidualization resulted in significant activation of many contractility-related gene ontologies (GOs), while conditioned medium reversed the trend. **(d)** GSEA plot showing negative enrichment of “Myotube differentiation” in dESFs in response to HTR8 conditioning; gene expression of leading-edge genes shown in heatmap. Key genes associated with contractile fiber maturation increased in dESF and reversed the trend in dEFS^cond^. **(e)** Many cardiac-related genes associated with actomyosin assemblage showed the same trend.

Further examination of a representative GO using Gene Set Enrichment Analysis (GSEA) showed that HTR8 conditioning resulted in negative enrichment for “Myotube differentiation”, with leading edge genes including MEF2C encoding Myocyte enhancer factor 2C, a key transcriptional regulator for myofibroblast transition, but also a key regulator of Ca^2+^ signaling associated with force generation in muscles [44]. We also found other genes encoding calcium channels which are crucial in triggering intracellular force generation, and actomyosin activation, particularly in the cardiomyocytes [45] **(Figure 3(d)**). These included RYR1, encoding Ryanodine receptor 1, a Ca^2+^ release channel in the endoplasmic reticulum, as well as KCNH1 encoding K^+^ voltage-gated channel [46]. We further calculated z-score for key genes associated with contractile fiber maturation in fibroblasts, and found them to be expectedly increasing during decidualization and reversing the trend in response to HTR8 conditioning. Again, many cardiac-related genes associated with actomyosin assemblage featured in the analysis shown in (**Figure 3(e)**). These included most importantly MYL9 encoding myosin light chain 9, the key homologue in fibroblast activation, which we have shown to be phosphorylated in dESF resulting in co-localization with actin filaments [39], ACTC1 encoding actin cardiac muscle 1, also present in airway smooth muscle cells [47], DMD which encodes dystrophin, a key gene in skeletal muscle fibers, TPM2 encoding tropomyosin 2, LDB3 which encodes a LIM domain binding cardiac protein, PARVA which encodes an actin-binding protein, as well as ACTN4 encoding actinin 4. Notably, actinin-4 has been shown to be critical for fibroblast contractility, while PARVA has been found to be upregulated in activated fibroblasts in pancreatic cancer [48]. Similarly, TPM2 has been found to be a prognostic marker in cancer-associated fibroblasts in colon carcinoma [49]. Overall, our data analysis confirmed the earlier findings that conditioned medium from HTR8 reduced the decidualization associated myofibroblast activation in ESFs.

### Computational segmentation enabled single-cell structural characterization

Processes such as cancer metastasis and placental development are fundamentally characterized by cellular spatial heterogeneity. However, evaluating the mechanical viscoelastic properties of tissues for individual single cells has been very challenging. We demonstrate our capability to virtually separate single cells and investigate the interactions between individual cells.

To achieve this objective, we indexed individual cells in an organoid and studied cell-cell structural interaction in the region of interest. **Figures 4(a-d)** show a zoom-in observation and analysis of cells representing a typical structural behavior in the ESF sample. **Figures 4(e-h)** show a typical pattern in the dESF sample. We can digitally separate cells of interest by hiding other cells (See **Figures 4(a-b)** and **4(e-f)**). **Figures 4(c)** and **(g)** show the index numbers given to the cells. **Figures 4(d)** and **(h)** show the von Mises strain, indicating the shear deformation. In the ESF sample, large strains were observed among fibroblast (red) cells. On the other hand, dESF sample shows the large strains among trophoblast (green) cells. This contrast between the two samples indicates that the decidualized fibroblast cells are more deformation-resistant than undifferentiated ESFs, which coincides with the well-known changes in decidualization, where endometrial fibroblasts (ESFs) undergo morphological and functional changes with increased cellular contractility in anticipation of implantation.

**Figure 4.**
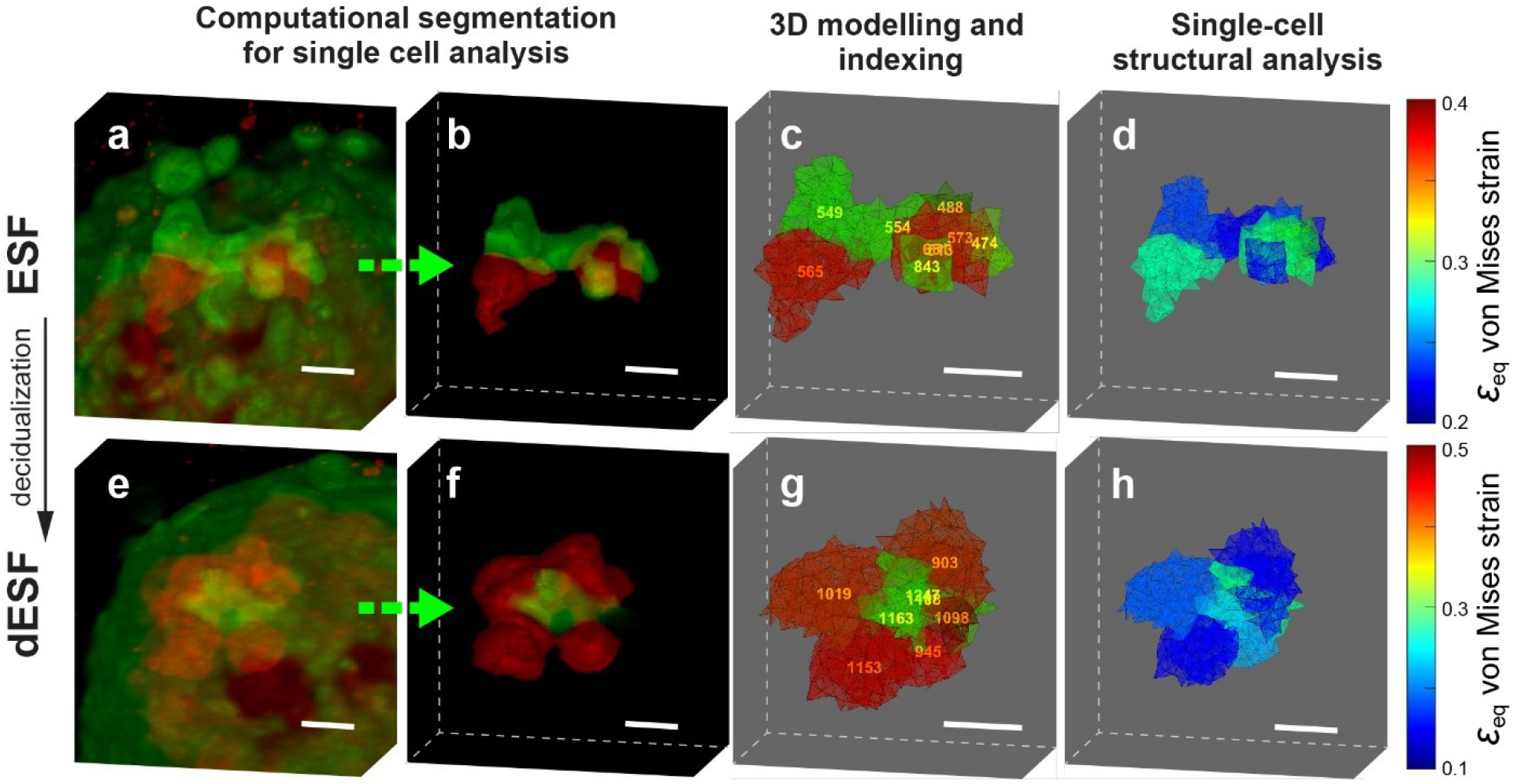
Computational single-cell analysis visualized a clear contrast in viscoelastic properties between **(a-d)** an ESF organoid and **(e-f)** a dESF organoid. We can highlight regions of interest by hiding other cells. In the ESF organoid, fibroblast cells (indicated in red, #513, #565, #573 and #651 in **(c)**) show the large strains in **(d)**. On the other hand, in the dESF organoid, trophoblast cells (indicated in green, #1108, #1163, and #1247 in **(g)**) experience the largest von Mises strains in **(h)**. These quantitative observations correspond well to the well-known structural changes in endometrial fibroblast decidualization.

### We combine single-cell visco-elastography and fluorescence cytometry to link molecular assay and 3D structural analysis

One innovative aspect of our method is its ability to correlate mechanical strain with the fluorescence of individual cells; the capability to study single cells allowed for both qualitative and statistical observation of viscoelastic transitions. In **Figure 5**, we show detailed single-cell analyses of the samples. For qualitative observation, we labeled examples of individual cells (#290,#336,#435, and #581 in ESF and #678, #701, #767, and #964 in dESF) showing increased fibroblast stiffness after decidualization. We then demonstrated a statistical analysis by creating unique 2D fluorescence-viscoelasticity correlation plots.

**Figure 5.**
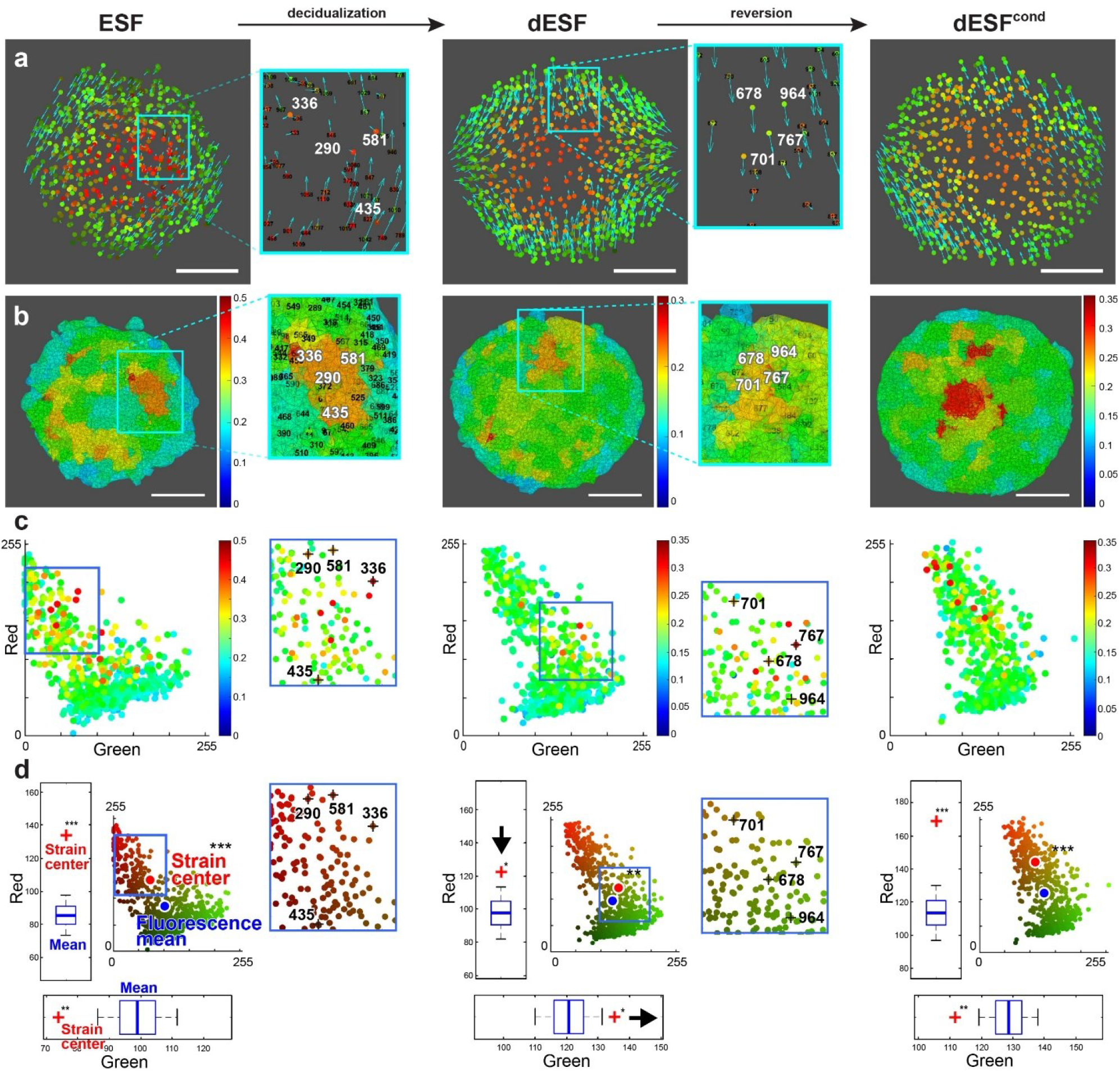
We conducted single-cell cytometry for qualitative observation and statistical analyses to demonstrate fibroblast viscoelasticity transition. In the ESF sample, larger strains are found among fibroblast (red) cells (example: #290,#336,#435, and #581), while those are found among trophoblast (green) cells (examples: #678, #701, #767, and #964) in the dESF sample. **(a)** Cellular displacement vectors in 3D organoids. Trophoblast cells (green, outer shell) tend to move in alignment with neighboring trophoblast cells in a solid-like motion. Fibroblast cells (red) in ESF and dESF^cond^ show fluid-like random motion, causing non-symmetric deformation. Eddy-like motion was observed in different scales. **(b)** Calculated von Mises strain to assess randomness in cellular motion. **(c)** von Mises stain plotted against 2D fluorescence. dESF showed larger strains in the trophoblast (green) region near fibroblasts (red). In ESF and dESF^cond^, larger strains were observed in the fibroblast (red) region near trophoblasts (green). **(d)** Statistical analysis of the strain distribution. The mean locations of the top 5% strain segments (strain center, large red points) are significantly different from the mean locations of randomly chosen 5% segments (fluorescence mean, large blue points) through two-dimensional Monte Carlo permutation tests (N = 10^9^). Both the green and red channels show clear reversed characteristics in dESF^cond^.

**Figure 5 (a)** shows the maps of internal cellular displacement. The vectors indicate the relative displacement with respect to the volume center of the organoid. We calculated the cellular displacement as the average displacement of the nodes contained in each cell. The measurement accuracy is high, with a deviation of 0.18 μm at an average of 29.6 μm displacement (see **Supplementary Information** and **Supplementary Figure 2(b)** for details). In EVTs (green), displacement vectors were overall well-aligned with those of neighbor trophoblast cells, showing characteristics commonly observed in a solid elastic material, which is expected for epithelial cells. In contrast, for ESFs (red), the displacement vectors showed a more liquid-like random spatial distribution. Remarkably, the decidualization of ESFs resulted in a dramatic change in the displacement vector map, with a more solid-like deformation pattern throughout the organoid samples. Treatment with the conditioned medium from HTR8 reverted the solid-like deformation pattern of dESFs towards a more random, liquid-like state. Through the von Mises strain analysis, we can quantify and visualize the regions with large strains (**Figure 5(b)**). As expected, dESF shows the smallest average strains. The calculated strains were significantly larger (p = 2.4 × 10^-24^) than the estimated fluctuation caused by the DIC tracking errors in nodal displacement (see the **Supplementary Information** for the error evaluation).

**Figure 5(c)** demonstrates the most notable capability of our approach, which allows a multifactorial assessment of a cell within its 3D tissue context, mapping the fluorescence of individual cells with a visualization similar to a flow cytometry plot. We observed that ESF and dESF^cond^ show larger strains in the regions with stronger red signals, indicating deformation in the fibroblast cells (red), while dESF shows larger strains in the regions with stronger green signals, indicating more deformation in EVTs (green). This comparison of the three conditions confirms the stiffening by decidualization and the reversed decidual stiffness after treatment with EVTs. The labeled cells shown in zoom-in images for ESF and dESF exemplify this structural transition through decidualization. In the ESF sample, cells #290,#336,#435, and #581, showing larger strains, are fibroblast (red) cells. In the differentiated dESF sample, cells #678, #701, #767, and #964, showing larger strains, are trophoblast (green) cells.

Our novel cytometry method also allows statistical correlation analysis of viscoelastic characteristics of individual cells with fluorescence information. In **Figure 5(d)**, we evaluated the relative statistical distance between the strain center and the fluorescence mean point. We used two-dimensional Monte Carlo permutation tests to find the mean locations of the segments with the top 5% strain (strain center, large red points) and the mean locations of randomly chosen 5% segments (fluorescence mean, large blue points). The permutation test for each condition (N = 109) showed that the strain center is significantly different from randomly chosen groups for all cases with p-values p = 5.0×10^-7^, p = 1.4×10^-3^, and p = 7.0×10^-7^, respectively. The statistical distances in the red and green axes were also separately calculated through permutation analyses and shown in boxplots in **Figure 5(d)**. The p-values are p = 3.2×10^-8^, p = 1.2×10^-2^, and p = 3.2×10^-7^ for ESF, dESF, and dESF^cond^, respectively, for the red channel, and p = 1.3×10^-3^, p = 1.8×10^-2^, and p = 3.2×10^-3^ for ESF, dESF, and dESF^cond^, respectively, for the green channel. The dESF organoid shows the strain center on the stronger side of the green signal and much closer to the fluoresce center in the red signal (indicated by the two arrows in the figure). The dESF^cond^ organoid shows reversed characteristics similar to ESF. The analysis demonstrated that the fibroblast cells increased the stiffness after decidualization, and the treatment with EVTs reversed the trend.

## Discussion

Identifying cellular viscoelastic transitions become the focus of recent studies [11-20]. However, it has been difficult for conventional microscopy-based assays to investigate the transitions of cellular and tissue structural characteristics, particularly in 3D and especially at heterogeneous tissue interfaces. Here, we demonstrated a method to quantify spatially-resolved characteristics of viscoelastic cell-cell interaction in 3D composite organoid models. Our method allowed statistical analysis of the viscoelastic properties of cells within the 3D microenvironment at the inter-tissue interfaces. We also demonstrated the capability of plotting the viscoelastic properties and multi-channel fluorescence signals, which could be combined with functional readouts, providing a method to create a multifactorial spatially resolved characterization of complex 3D microtissue structures.

We applied our method to study the dynamic physiology of the maternal-fetal interface in humans. Our 3D analysis confirmed existing beliefs, discovered new insights, and validated recent findings regarding the co-adaptation in the evolution of great apes. We showed that decidualization of endometrial stromal fibroblasts (ESFs) dramatically increases tissue stiffness, acquiring a solid-like characteristic. We also confirmed in 3D that the interaction with fetal trophoblasts can reverse this trend, rendering dESFs more liquid-like and likely more invasive. Indeed, the RNAseq analysis of dESF^cond^ revealed a reversal in numerous gene sets linked to actomyosin contractility and force generation. The invasion of placental trophoblasts into the endometrial stroma occurs in all hemochorial species, with great apes displaying particularly aggressive behavior. Our method demonstrates that these mechanisms could be explained partly by the changed viscoelastic properties of the decidual compartment at the maternal-fetal interface.

There are many parallels between placental invasion and epithelia-stroma interactions in other essential biological events. The formation of organs, including folding of dermal layers, wrinkling of dermis, keloids and scar formation, or epithelial-stroma invasion, is guided by the differences in neighboring tissue mechanics. The centrality of tissue mechanical properties in defining cell specification, development, morphogenesis, and pathogeneses of various kinds has been well-established [18-20]. Our method can potentially provide a mechanistic explanation centered on the mechanics of tissues underlying the dysregulation of interfaces in various contexts. One could easily use our method to determine cell-surface markers or signaling readouts in the 3D milieu. Hence, our tool broadly expands the applicability of biomedical imaging to tissue characterization, including phenotyping tumor metastasis in cancerous tissues, studying embryonic body development, and assessing wound healing and regeneration.

Our study is among the first to apply the digital image correlation (DIC) technique to high-end 3D microscopy for tissue viscoelastic characterization. Structural analysis is based directly on processing fluorescence images, revealing the link between molecular and structural studies. DIC computes sample shape, motion, and deformation information through the feature analysis of digital images. With the recent rapid advancement in digital photography and microcopy, it is gaining more attention as a technique for materials characterization. In our structural analysis, the resolution and the sample size limit are mainly determined by those of the image acquisition method. Spatial-domain interpolation or frequency domain analysis allows DIC analysis to obtain resolution beyond digital image pixel resolution. Using a high-end commercial light-sheet microscope, we measured cellular displacement beyond the optical resolution limit (see **Supplementary Information** for details). For further resolution improvement, our method can be flexibly applied to various types of imaging modalities. Recently reported light-sheet microscopy techniques have demonstrated optical resolution close to ∼100 nm [26,50]. Our method can potentially study the intracellular structural characteristics of individual cells, which can be spatially correlated to the fluorescence from molecular markers.

In conclusion, our novel method is a powerful yet flexible tool for characterizing cellular-scale viscoelastic properties in 3D models, with broad applicability. As fluorescence microscopy is now a crucial tool in quantifying cellular signaling, our method will link mechanical and structural properties with the intracellular signaling state of cells in heterogeneous 3D ensembles.

## Acknowledgement

The authors acknowledge NSF (CAREER - 1942518 and IIBR - 2223957) and NIH (NCI - R37CA248161) for funding.

