## Supplementary Information for "Lightsheet microscopy integrates single-cell optical visco-elastography and fluorescence cytometry of 3D live tissues"

#### 1. Creating 3D models

**Suppl. Fig. 1** shows the 3D modeling procedures. The initial 3D image before indentation (i.e., the first timeframe) was used to create the model. **Suppl. Fig. 1(a)** shows a slice of a raw 3D volume, which typically consists of ~360 z layers of 1600 × 1600 pixel xy images. We then Gaussian 3D low-pass and high-pass filtered the volume to remove the background variations and small noises. The cut-off frequencies were chosen for the filter bandwidth to include the typical cell sizes (5-15 μm). Cells were found as bright spots after the filtering (**Suppl. Fig. 1(b)**). We used a MATLAB function 'watershed' to separate these cells into 3D segments (**Suppl. Fig. 1(c)**). This 'watershed' based segmentation process is modified from [1]. For strain analysis, we used the hemispherical region closer to the microscope objective to have a better imaging quality. Each segment is numbered for further single cellular analysis. The number of segments in the region was typically ~500-600. We then evenly distributed nodal points on the segment boundaries and within segments, keeping distances of ~10 pixels (~ 4μm) between. These nodal points were used for 4D digital image correlation (DIC) and for creating tetrahedral mesh using the MATLAB 'alphaShape' function. The total number of tetrahedral elements and nodal points studied are typically ~20,000 and ~100,000-200,000, respectively. Each segment contained ~100-200 tetrahedral elements and ~50 nodal points. The number of tetrahedral elements is more than that of the nodes because a node is usually shared by multiple elements. Nodes on segment boundaries are shared by multiple segments. Each element only belongs to one segment that contains it.

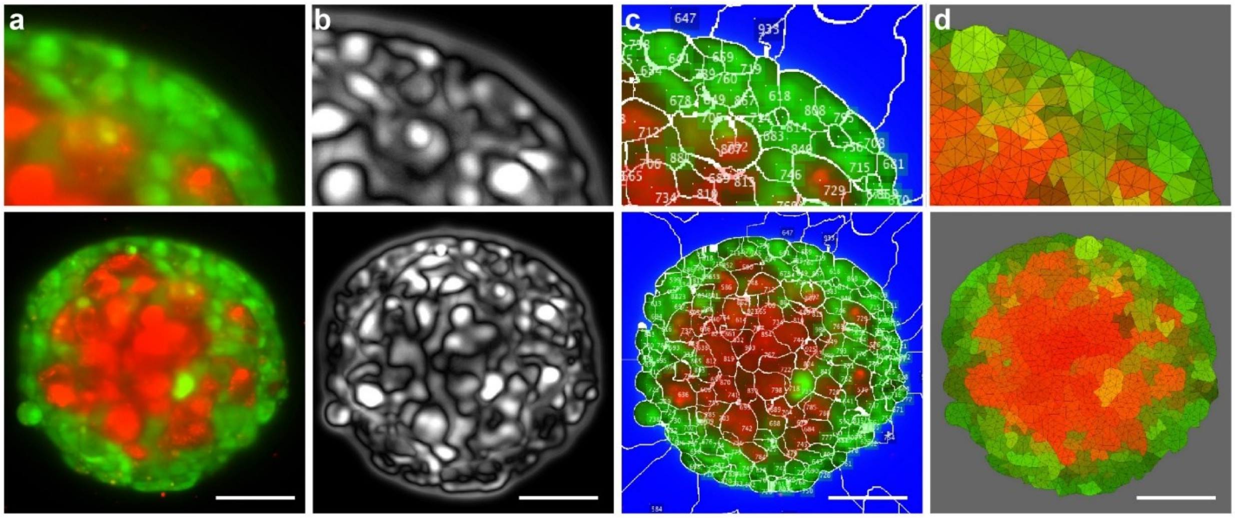


**Supplementary Figure 1.** Steps of creating a 3D model. **(a)** A slice of the raw 3D image. **(b)** The 3D volume is low-pass and high-pass filtered to find cell features. **(c)** The 3D watershed technique is used to divide the 3D volume into ~500-600 segments representing single cells. Each segment is numbered for further analysis. **(d)** We distributed a total of ~20,000 nodal points for meshing and optical tracking (digital image correlation). Each segment is meshed to 100-200 tetrahedral elements. Scale bars = 50 μm.

### 2. Displacement and strain analysis through 4D digital image correlation

The digital image correlation (DIC) analysis was based on spatial-domain volumetric pattern matching. The volumetric region of 30 pixels × 30 pixels × 14 pixels (= 11.6 µm × 11.6 µm × 7.0 µm) around each nodal point was chosen and tracked along the indentation steps. Using interpolation, we found the displacement in real numbers beyond digitized pixel resolution given in integers.

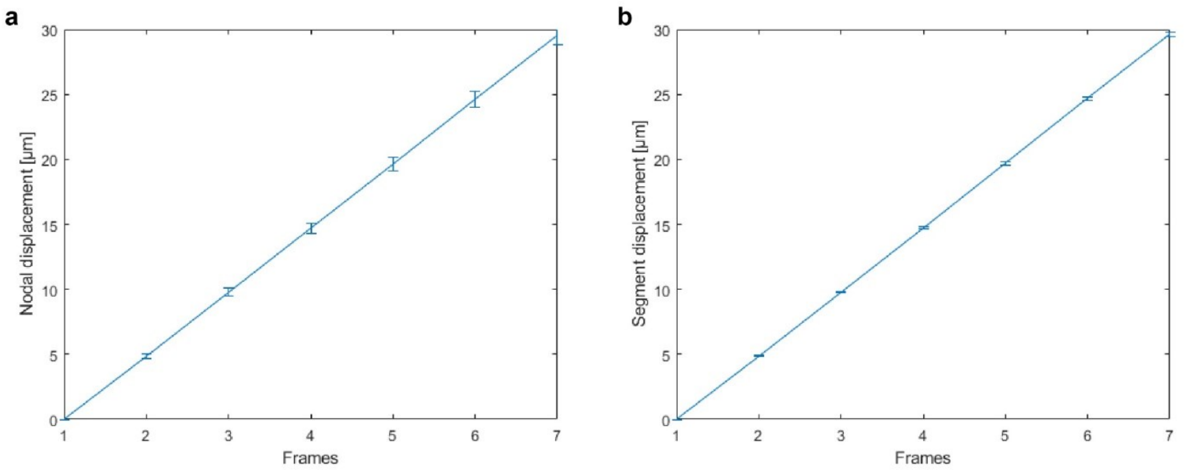


**Supplementary Figure 2.** Evaluation of the digital image correlation (DIC) accuracy. An organoid sample was given 6 steps (total 7 timeframes including the initial image) of 5 µm translation in x direction without compression, and the displacement was measured by our 4D DIC algorithm. **(a)** Nodal displacement of 9638 nodes. The deviation from the linear fit ($R^2$ = 0.99999) was 0.73 µm at 29.6 µm indentation. **(b)** Segmental displacement measured for 385 segments. The deviation from the linear fit ($R^2$ = 0.99999) was 0.18 µm at 29.6 µm indentation, which was smaller than the optical resolution defined by the Rayleigh criterion (0.61 × λ) /NA for the fluorescence wavelengths (517 nm and 602 nm).

The accuracy of 4D digital image correlation (DIC) was evaluated by tracking an organoid given a translational movement without any compression or stretching. **Suppl. Fig 2(a)** shows the measured displacement of 9638 nodes for 6 indentation steps (R = 0.99999). The average displacement after 6 steps was 29.6±0.73 µm (127.6± 3.2 pixels). We calculated the cellular displacement as the average displacement of all the nodes in each segment. Supplementary **Suppl. Fig 2(b)** shows the displacement measurement for the same translational movement (R = 0.99999). The average displacement after 6 steps was 29.6±0.18 µm (127.6± 0.8 pixels). For comparison, we note that the microscope optical resolutions defined by the Rayleigh criterion (0.61 × λ) / (N.A.) for the fluorescence wavelengths of 517 nm and 602 nm are 0.32 µm and 0.37 µm, respectively. The errors in nodal displacement caused noise in the von Mises strain calculation. We observed a noise component of 0.18±0.07 (N = 385) at an average translational displacement of 29.6 µm, while a compression testing with a comparable 30.4 µm average displacement showed an average von Mises strain of 0.22±0.09 (N = 631), which was significantly larger than the translation ($p = 7.0 \times 10^{-16}$).

#### 3. Sample preparation

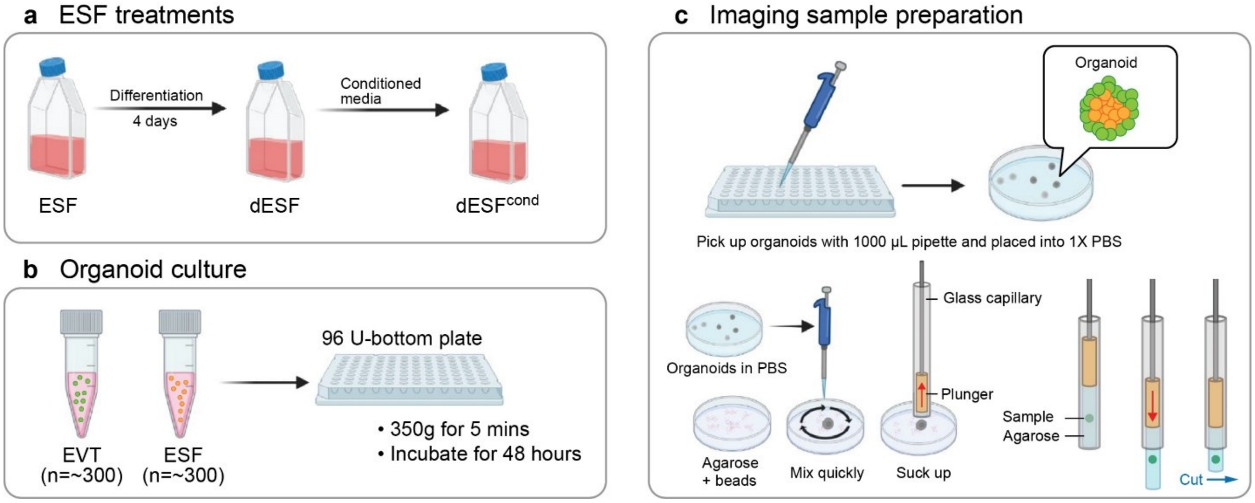


**Supplementary Figure 3.** **(a)** We prepared four types of ESFs, namely, ESF, dESF, and dESF$^{cond}$. **(b)** Organoids were prepared by counting 300 trophoblast and 300 ESFs using a hemocytometer and mixing them in a 96 U-bottom well plate. **(c)** Samples for imaging were embedded in agarose suspended from a glass capillary.

**Suppl. Fig. 3** summarizes the protocol of organoid culture and imaging sample preparation. The details are described as follows:

**Cell treatments.** We prepared organoids containing human endometrial stromal fibroblasts (ESFs) and placental extravillous trophoblast cells (EVTs). ESFs were derived from primary cells isolated from a normal patient at the authors' institution. For EVTs, we used HTR8-SV-Neo (HTR8), a cell line representing extravillous trophoblasts with partial mesenchymal properties, purchased from ATCC (CRL-3271). ESFs were maintained in DMEM/Hams F-12 50/50 Mix (phenol-red free), supplemented with 10% charcoal-stripped fetal bovine serum (CS-FBS) and 1X antibiotic-antimycotic. Decidualized ESFs (dESFs) were prepared by treating ESFs at 80% confluency with 0.5 mM 8-B-cAMP and 1 mM medroxyprogesterone acetate (MPA) for four days in DMEM/F12 (with phenol-red) supplemented with 2% fetal bovine serum (FBS). We refer to this medium used for cell differentiation as "differentiation media." Conditioned dESFs (dESF$^{cond}$) were prepared by treating pre-decidualized dESFs for 2 days with conditioned medium from HTR8 cells. To obtain the conditioned media, we cultured HTR8 cells in SIGMA RPMI-1640 supplemented with 10% (FBS) and 1 X antibiotic-antimycotic. The conditioned media from the HTR8 flask was collected after a minimum of two days of growth, diluted 1:1 with differentiation media, and then placed onto dESF cells for two days.

**3D organoid culture.** After the completion of ESF treatment immediately before 3D culture, cells were labeled with cell tracker dyes. HTR8 cells were labeled with CellTracker™ Green CMFDA (5-chloromethylfluorescein diacetate) (Thermo Fisher Scientific, Waltham, MA) at a 1:1000 ratio, diluted in 1x phosphate-buffered saline (PBS) for 14 minutes in a cell incubator while being gently shaken every 5 minutes. Similarly, the endometrial fibroblast cells were labeled with Invitrogen by CellTracker™ Red CMTPX (Thermo Fisher Scientific, Waltham, MA) at a 1:500 dilution in 1x PBS for 35 minutes, with gentle shaking every 5 minutes. To confirm the efficacy of the labeling process, we observed the cells under an inverted fluorescent microscope. The cells were then washed three times with 1X PBS and detached using Gibco 0.25% trypsin-EDTA.

We prepared the organoids by counting and mixing EVTs and ESFs using a hemocytometer. The cells were seeded into a U-bottom plate, followed by centrifugation for 5 minutes at a speed of 350 g. The cell mixture was then cultured in a solution where 1X of methyl cellulose was diluted in differentiation media. We excluded 8-B-cAMP due to its toxicity to the HTR8 cells. Through multiple experiments, we have determined the optimal size for imaging was an organoid seeded from 300 trophoblast and 300 ESFs. The plate was kept in a humidified incubator at 37ºC, adhering to a 48-hour incubation period. Deviating beyond this time frame could compromise the quality of the fluorescent signal, which is integral to our analysis. We found that HTR8 and ESFs spontaneously sorted themselves spatially in 3D, with the ESFs in the core, surrounded by layers of HTR8, effectively creating a placenta-decidua interface. We carefully aspirated organoids from the U-bottom plate using a 1000 pipette and transferred them into a 35 mm by 10 mm petri dish containing 1X PBS. We picked up samples that were between 160-180 µm in diameter.

**Preparation of samples for imaging.** For 3D imaging, organoids were embedded in low melting point agarose gel (MilliporeSigma, Burlington, MA). Agarose gel prepared in a 250 mL glass bottle at 0.8% concentration was placed on a 250ºC hot plate until it became liquified and clear. We transferred 2000 µL of heated agarose into an empty 35 mm × 10 mm petri dish using a 1000 µL Eppendorf micropipette. We then pipetted 3 µL of Invitrogen™ FluoSpheres™ Carboxylate-Modified Microspheres, 0.2 µm (Crimson) (Thermo Fisher Scientific, Waltham, MA) into the 2mL agarose and mixed using the micropipette tip. The beads were sonicated for 10+ minutes before use. We used a Fluke 62 Max+ Infrared Handheld Thermometer to monitor the agarose temperature and waited until it cooled down to 40ºC before adding the organoid. A 1000 µL Eppendorf micropipette set to 100 µL was used to pick up organoids from the petri dish. The organoids with the PBS media were mixed into the cooled melted agarose. We used a Wiretrol® II 50 & 100 µL glass capillary (I.D. 1.4 mm) with a wire plunger (Drummond Scientific, Broomall, PA) to suck up the organoid containing agarose. The position of the plunger is manually adjusted to have the organoid at a middle point of the agarose in the capillary. Once the organoid position is fixed, we let the agarose cool down to 26ºC. When the gel is cured, a scalpel is used to cut off the excess agarose, ensuring a clean, straight cut at the end. The sample is now ready for light-sheet imaging. To prevent dehydration of agarose during transfer, we kept the capillary tip containing the sample in a 15 mL conical tube containing PBS.

#### 4. RNA sequencing and Gene Set Enrichment Analysis

Cells were lysed and RNA was isolated using RNeasy Mini Kit (Qiagen). Bioanalyzer 2100 (Agilent) was used to evaluate the integrity of RNA, and samples with RIN ~8 were processed further for library preparation. Novogene Inc. performed the library prep and RNA sequencing. NCBI GRCh38 genome assembly was used to align RNA reads using HISAT2 pipeline. HTSeq was used to count the reads [2], and DESeq2 was used to estimate p-values and fold-changes for differential expression between undifferentiated, decidualized, and conditioned samples [3].

Moderated log2fold change was used for differential analysis. P-values for differential expression were calculated using the Wald test. For gene ontology (G.O.) or pathway (Kegg) activation analyses, the Fisher exact test was used to calculate the overpresentation of terms using a hypergeometric test followed by multiple testing corrections [4]. Transcription factor (T.F.) activation scores were calculated using Ingenuity Pathway Analysis (IPA, Qiagen Inc.). For Gene Set Enrichment Analysis (GSEA), we used methods developed by Subramanian et al. [5].
